# Comparative transcriptomics reveals the molecular mechanisms of maize in response to downy mildew caused by *Peronosclerospora philippinensis* (Weston) Shaw

**DOI:** 10.1101/2023.11.05.565721

**Authors:** Edward Cedrick J. Fernandez, Eliza Vie M. Simon-Ada, Jay-Vee S. Mendoza, Anand Noel C. Manohar, Roanne R. Gardoce, Tonette P. Laude, Fe M. Dela Cueva, Darlon V. Lantican

## Abstract

Maize is the Philippines’ second most valuable staple crop based on overall value and total area planted. Yellow maize, which is the most important type, is utilized as feed for poultry and swine. Still, local maize production is hampered by the Philippine downy mildew (DM) disease caused by the pathogen *Peronosclerospora philippinensis* (Weston) Shaw, causing substantial losses in maize production annually. To comprehend the underlying resistance mechanisms upon the presence of the disease, we performed an RNA-Seq comparative transcriptomics approach between mock-inoculated and DM-inoculated susceptible and resistant yellow maize inbred lines. Among the identified differentially expressed genes (DEGs), we detected 43 DEGs shared in both genotypes which may play roles in the basal defense response of maize upon DM infection. We also identified 68 DEGs exclusive to the susceptible genotype, providing insights into the molecular responses underlying successful DM disease progression in maize. Further, we detected 651 DEGs unique to the resistant genotype. This set of genes revealed that the molecular basis of DM resistance in maize is governed by multi-faceted defense strategies. These include multi-process regulations such as programmed cell death (PCD), regulatory proteins, and transcription factors involved in pathogen defense mechanisms, cell wall organization, homeostasis, and many others. Out of 694 resistant DEGs detected, we narrowed down genes of interest, particularly those highly overexpressed and associated with disease resistance found in other crops. Finally, transcriptome-wide variants and their corresponding impact on gene function were detected for further application in targeted genotyping-by-sequencing, association studies, and marker-assisted DM resistance breeding.

## INTRODUCTION

Maize (*Zea mays* L.) is the second most valuable crop in the Philippines, based on overall value and total area planted (De los Santos et al. 2007). The country’s two major types of maize, yellow and white, have distinct utilization (Salazar et al. 2021). White flint maize is mainly used as a substitute staple for rice whereas yellow maize is a major component of the animal feed industry. Yellow maize demand and productivity continue to soar due to the increasing demand for meat consumption. Based on the latest crop statistics report in 2021, maize production grew to 8.30 from 7.91 million metric tons, 73% of which is devoted to yellow maize production (PSA 2023). As both maize demand and productivity soar, problems linger due to different biotic and abiotic factors including the continuous devastation brought about by crop diseases such as the downy mildew disease.

The Philippine downy mildew (DM) caused by the pathogen *Peronosclerospora philippinensis* (Weston) Shaw, is one of the major pathogens in the country (Dela Cueva et al. 2022) and is considered the most virulent of downy mildew pathogens in maize (Murray 2009). *P. philippinensis* bears a huge resemblance to *Peronosclerospora sacchari*, the causal pathogen of downy mildew in sugarcane, in terms of morphology and symptoms. A 40-60% yield reduction is expected under normal conditions, however, favorable conditions for disease progression can lead to 80% to complete annihilation (Exconde and Raymundo 1974). Deployment of resistant cultivars remains the most preferred means for disease control, including against downy mildew (Lukman et al. 2013). Inbred line Ki3 was reported to be highly resistant in the Philippines (disease incidence of 0-1%; George et al. 2003) while a few DM-resistant maize breeding materials such as Nei 9008 and P345, have already been developed in the country (Raymundo and Exconde 1976; Pascual et al. 2006). Recently, Ada et al. (2023) reported that maize inbred line CML 431 exhibited downy mildew disease resistance in response to *P. philippinensis*, corroborating the result of the inbred line’s resistance to downy mildew disease caused by *P. sorghi* conducted by CIMMYT (2015). Meanwhile, the use of the systemic fungicide metalaxyl has effectively controlled losses attributed to the disease (Anahosur and Patil 1980). However, previous reports have already suggested that the effectiveness of metalaxyl both as seed treatment and fungicide has already declined (Lukman et al. 2013; Ginting et al. 2020). Hence, it is imperative to initiate breeding programs aimed at utilizing DM-resistant lines (Fernandez et al. 2023) to effectively address the long-term devastating effects of DM in the country.

Understanding the plant-pathogen interactions is a precursor to the design and implementation of strategies for minimizing the economic shortages caused by the pathogen on crop production (Boyd et al. 2013). Although we have existing breeding materials with sources of resistance, the nature of the interaction between maize and downy mildew needs to be unraveled at the molecular level. Comparative transcriptome analyses through next-generation sequencing (NGS) technologies have elucidated the mechanisms of disease response in many different crops (Lantican et al. 2023). For instance, mechanisms of maize resistance for the gray leaf spot (GLS) caused by the fungi *Cercospora zeae-maydis* and *Cercospora zeina* were studied by this approach (Yu et al. 2018). More recently, a transcriptome meta-analysis using publicly available RNA-seq data sets from seven independent maize fungal pathogen inoculation experiments was performed to elucidate the molecular mechanisms of resistance against multiple disease-causing fungi in maize (Wang et al. 2022). Nevertheless, there are still no reports of the same approach explaining the mechanisms of maize-DM interactions in terms of resistance response and/or disease progression. Here, we give insights into the molecular basis of disease resistance between DM-susceptible and DM-resistant yellow maize inbred lines in response to the onset of DM infection by analyzing the set of DEGs between a susceptible and resistant maize line as well as identifying transcriptome-wide variants (SNPs and indels) based on the generated RNA-seq data.

## MATERIALS AND METHODS

### Preparation of samples and DM inoculation

DM-susceptible yellow maize line Philippine inbred Pi23 and DM-resistant CML 431 maintained at the Institute of Plant Breeding, University of the Philippines Los Ban os (IPB-UPLB) were used for RNA-Seq-based transcriptomics analysis. The Philippine inbred Pi23 is an elite yellow maize inbred developed by IPB-UPLB which is used as a parental line of a released single-cross hybrid susceptible to DM (Ruswandi et al. 2014) while CML 431 is another yellow maize inbred from the International Maize and Wheat Improvement Center (CIMMYT) with reported resistance against DM (Ada et al. 2023; CIMMYT 2015). Three (3) biological replicates, each for mock- and DM-inoculation, for both resistant and susceptible lines for a total of 12 biological samples were sown and grown in large pots filled with sterilized soil and kept inside separate 50-mesh cages in a greenhouse at IPB-UPLB.

Infection was administered via the whorl inoculation method following conidial spore collection (Singh and Gopinath 1985). Leaves from the DM-infected sugarcane plants were sourced, covered with wet polyethylene bags, and incubated overnight at 24 °C and 80% humidity to facilitate conidial formation. Spores were collected at 4 AM (GMT +8) by brushing them into distilled water followed by transferring them to a sterile container. The spore suspension was quantified and adjusted to 10^6^ CFU/ml. Plants were inoculated by dropping 1 ml of spore suspension inside the whorl. As for the mock-inoculated samples, 1 ml of sterile distilled water was dropped for each plant. Disease progression was validated visually via the presence of DM symptoms i.e., chlorosis/streaking in the leaves. Finally, leaves were collected from the youngest leaf tissues of maize plants 72 hours post-inoculation (hpi) for RNA extraction.

### RNA extraction of resistant and susceptible yellow maize lines

Total RNA was extracted from the youngest leaf tissues of three biological replicates of DM-susceptible and DM-resistant yellow maize inbred lines (mock-inoculated and DM-inoculated) using the modified NucleoSpin™ Mini Kit for RNA purification (Macherey-Nagel™, GmbH & Co. Du ren, Germany). One hundred fifty (150) mg of plant tissues from each sample were homogenized in liquid nitrogen in a double sterilized mortar and pestle.

RNA concentration and quality were determined by gel electrophoresis in 1% UltraPure™ agarose (Invitrogen Corp. Carlsbad, California, USA) in 1× TBE running buffer at 100 V for 40 min. The gel was stained with 0.1 uL Gel Red (Biotium, CA, USA) visualized under UV illumination at 300 nm using the Enduro GDS Touch Imaging System (Labnet International, Inc, Edison, New Jersey, USA).

### Next-generation sequencing (NGS) of RNA-extracted materials

The extracted high-quality RNA samples from both control and DM-infected susceptible and resistant yellow maize inbred lines were sent to the Philippine Genome Center – DNA Sequencing Core Facility (PGC-DSCF), University of the Philippines Diliman, Diliman, Quezon City, Philippines for outsourced next-generation sequencing. Quality control check was done through RNA quantitation, 260/280 and 260/230 nm absorbance (OD) ratio measurements, and gel separation using Microchip Electrophoresis System (MCE™-202 MultiNA; Shimadzu Biotech, Kyoto, Japan). Sequencing libraries were constructed using TruSeq Stranded Total RNA Library Prep Plant (Illumina), followed by next-generation sequencing on the Illumina NextSeq 500/550 sequencing platform.

### Bioinformatics analyses

The bioinformatics pipeline for RNA-seq transcriptomics analysis utilized in the study is presented below (Figure 1). The paired-end raw transcriptomic sequences were pre-processed by removing the adapter sequences and low-quality base score nucleotide sequences using *Trimmomatic* v0.36 (Bolger et al. 2014) with default parameters except: SLIDINGWINDOW:5:15. The qualities of the raw and trimmed reads were assessed by *FastQC* v0.11.8 (Andrews 2010). The paired-end reads were mapped to the *Zea mays* L. version 5 B73 genome (Woodhouse et al. 2021) using *STAR* v2.7.10b (Dobin et al. 2013) default settings. The gene expression abundances were estimated from the resulting binary alignment map (BAM) files using RNA-Seq by Expectation-Maximization (RSEM; Li and Dewey 2011) at default settings. A matrix of the read count data generated from RSEM quantification was prepared and imported to R package *DESeq2* v1.26.0 (Love et al. 2014) using *tximport* v1.20.0 (Love et al. 2017).

**Figure 1.**
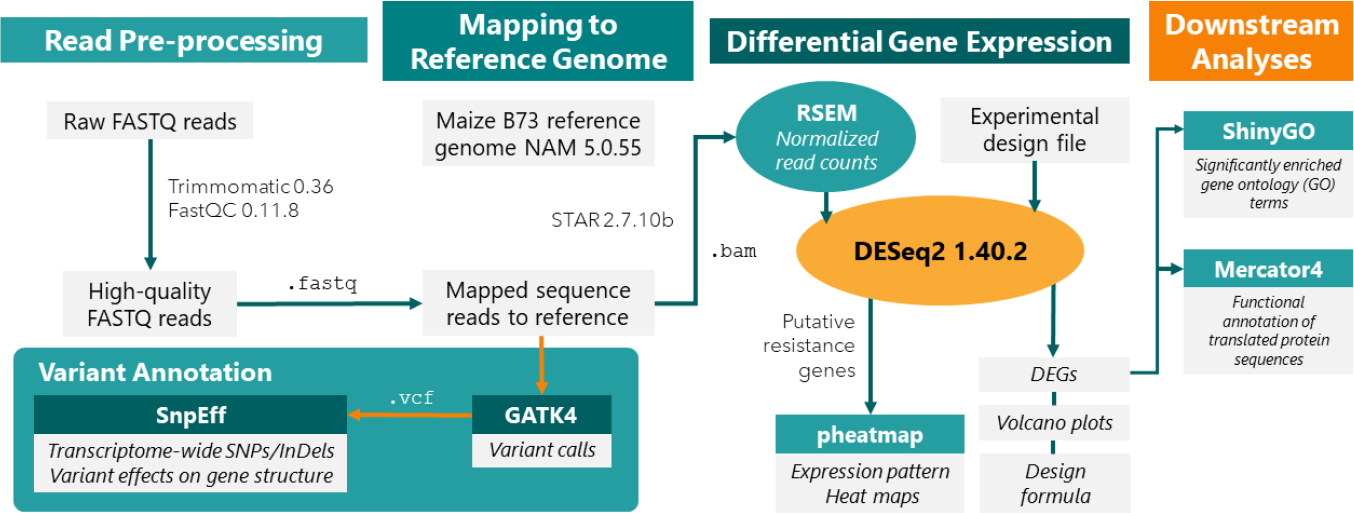
Schematic diagram of the bioinformatics pipeline implemented to compare the transcriptomic profiles of DM-resistant CML 431 and DM-susceptible Pi23, in response to DM infection using mock-inoculated samples as baseline control.

A single-factor design was implemented independently from each inbred line to consider the factor introduced by the differences in the inoculation state from biological replicates. The relative log normalization (RLE) method determined the DEGs between the mock-inoculated and DM-inoculated samples. P-values were determined using the Wald test from the general linear model (GLM) fitting. The Benjamini-Hochberg (BH) *p*-value adjustment procedure was used to calculate the false discovery rate (FDR) values. Genes having FDR values of <0.05 were deemed differentially expressed between mock-inoculated and DM-inoculated susceptible Pi23 and resistant CML 431. Representative heat maps for DEGs to visually show differential expression patterns were generated using the R package *pheatmap* v1.0.12 (Kolde 2012).

Gene ontology (GO) enrichment analysis was performed using ShinyGO v.0.77 (Ge et al. 2020) where significantly enriched GO terms found in the resistant and susceptible maize lines were analyzed with an FDR value of 0.05. Also, the translated proteins from the DEGs were extracted and uploaded to Mercator4 release 5 (Bolger et al. 2021) to assign functional annotations to the protein sequences. Finally, an exploratory analysis was performed to screen for DEGs conferring disease resistance such as genes for pattern recognition receptors (PRRs), nutrient transporter genes, regulatory proteins, transcription factors, etc.

Pre-processing, variant calling, and variant annotation were also performed after alignment to the reference genome using the Genome Analysis Toolkit (GATK) best practices recommendations for RNA-seq to detect high-confidence variants related to DM-resistance in maize (Van der Auwera and O’Connor 2020). Exclusive and shared transcriptome-wide variants (i.e., SNPs and indels) between the infected susceptible and resistant maize lines were analyzed using BCFTools v.1.9 (Danecek et al. 2021). Finally, the filtered SNPs and indels were functionally annotated using SnpEff v.5.1 to predict their effects on gene structures (Cingolani et al. 2012).

## RESULTS

### RNA-seq transcriptome data analyses of yellow maize inbred lines in response to DM infection

To detect important genes that may play roles in Philippine downy mildew (DM) resistance in maize, a comparative RNA-Seq transcriptomics analysis was performed in two yellow maize inbred lines, specifically, CML 431 (DM-resistant; Ada et al. 2023) and Pi23 (DM-susceptible; Ruswandi et al. 2014). The inbred lines were subjected to mock and DM inoculations using the *P. philippinensis* reference isolate from the Plant Pathology Laboratory-Institute of Plant Breeding (Aguilar et al. 2022). The RNA samples were isolated from young leaf tissues at 72 hours post-inoculation (hpi). A total of 12 RNA samples were sent for outsourced next-generation sequencing using Illumina NextSeq 500/550 to represent the total transcriptomic profiles of control (mock-inoculated) and DM-inoculated resistant and susceptible yellow maize inbred lines. Approximately 30 to 46 million raw reads (35-151 bp length per read) were generated (Supplementary Table S1).

The maize B73 reference genome assembly version 5 (Zm-B73-REFERENCE-NAM-5.0.55) retrieved from the MaizeGDB (Woodhouse et al. 2021; https://www.maizegdb.org/) was used as a reference in the genome-guided mapping of the RNA-seq reads for differential gene expression transcriptomic analysis. Approximately 9 to 18 million trimmed paired-end reads (∼250-bp length per spot) were observed to uniquely map to the reference genome. On average, 82.12% (mock-inoculated ‘Pi23’), 81.71% (DM-inoculated ‘Pi23’), 76.09% (mock-inoculated ‘CML 431’), and 82.90% (DM-inoculated ‘CML 431’) were mapped to the genome (Supplementary Table S1).

### Expression patterns between inbred lines Pi23 (susceptible) and CML 431 (resistant) in response to DM

The RSEM-normalized expected count data from resistant DM-infected CML 431 (27,036 genes with non-zero read count) and susceptible Pi23 (27,782) were statistically compared using the DESeq2 R package (Love et al. 2014). The DESeq2 pipeline was able to identify 694 and 111 differentially expressed genes upon DM infection in CML 431 and Pi23, respectively (Supplementary Table S2). Of these significantly expressed genes, 25 were up-regulated while 86 were down-regulated in Pi23. Meanwhile, 248 genes were up-regulated while 446 genes were down-regulated in the DM-resistant CML 431 (Volcano plots; Figure 2). Moreover, a comparison of the DEG profiles between the susceptible and resistant yellow maize inbred lines revealed 651 DEGs detected only on the resistant inbred CML 431 (Venn Diagram; Figure 2), while 68 exclusive DEGs were detected in the susceptible inbred Pi23. This distinct set of genes exclusive to Pi23 offers valuable insights into the molecular responses underlying the successful DM disease progression in maize. Moreover, a similar expression profile was observed for 43 DEGs in both inbred lines, indicating the shared roles of these genes in the basal response of maize to DM infection, irrespective of their resistance or susceptibility response.

**Figure 2.**
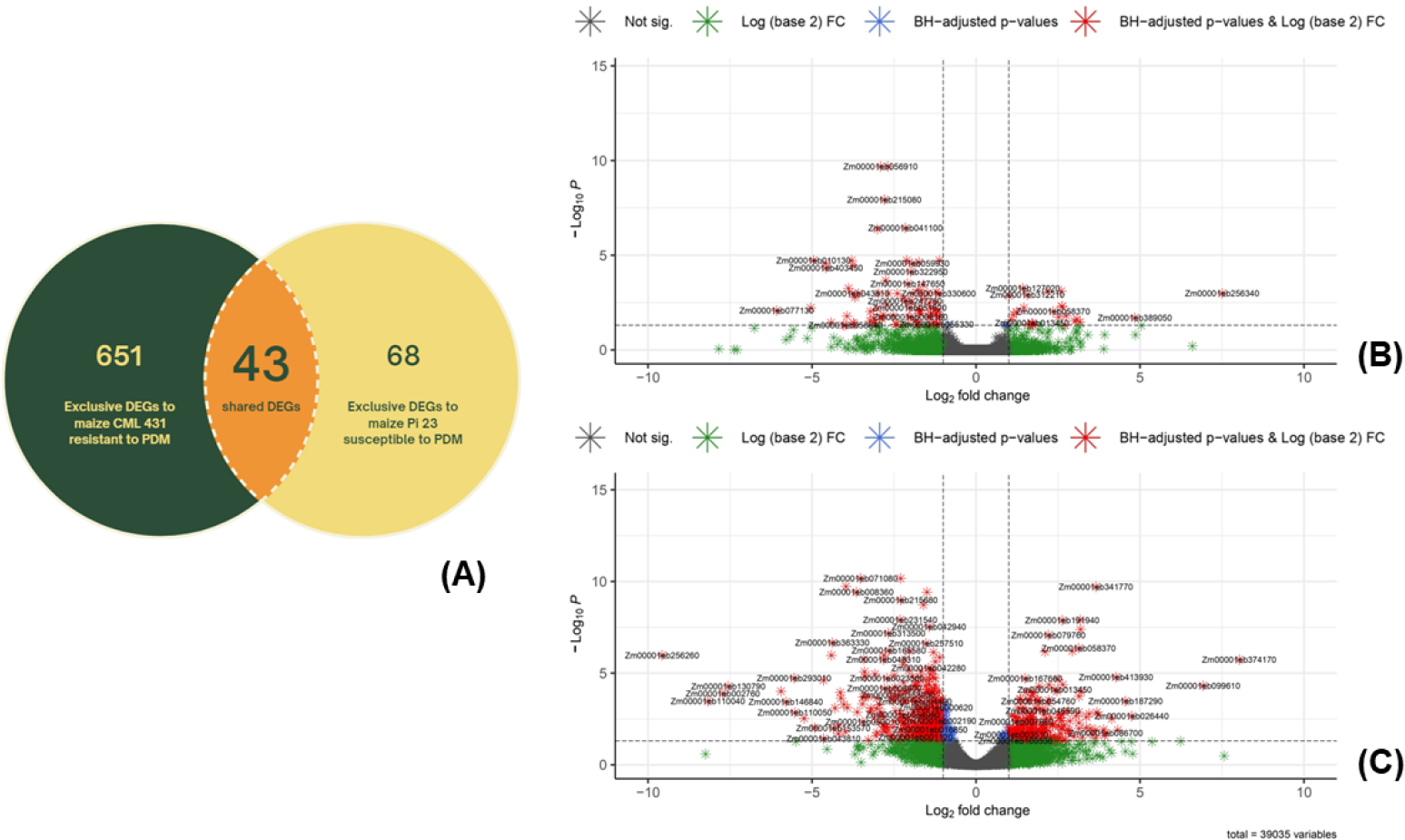
Volcano plots and Venn diagram of DEGs. (**A**) Venn diagram showing the overlap of DEGs between DM-susceptible Pi23 and DM-resistant CML 431. Volcano plots of the differentially expressed genes (DEGs) in susceptible Pi23 (**B**) and resistant CML 431 (**C**) three (3) days post-inoculation. The x-axis shows the fold change difference in the expression of genes, and the y-axis indicates the BH-adjusted p-values for the changes in the expression. An absolute value of log2 fold change >1 and the *p*-value < 0.05 was set to declare differentially expressed genes (DEGs).

### Gene ontology (GO) enrichment of DEGs and gene product functional annotation

To understand the gene and gene product functions of the DEGs in response to DM, the significantly enriched GO terms and functional annotation of the gene products were assessed using ShinyGO (Ge et al. 2020; Figure 3).

**Figure 3.**
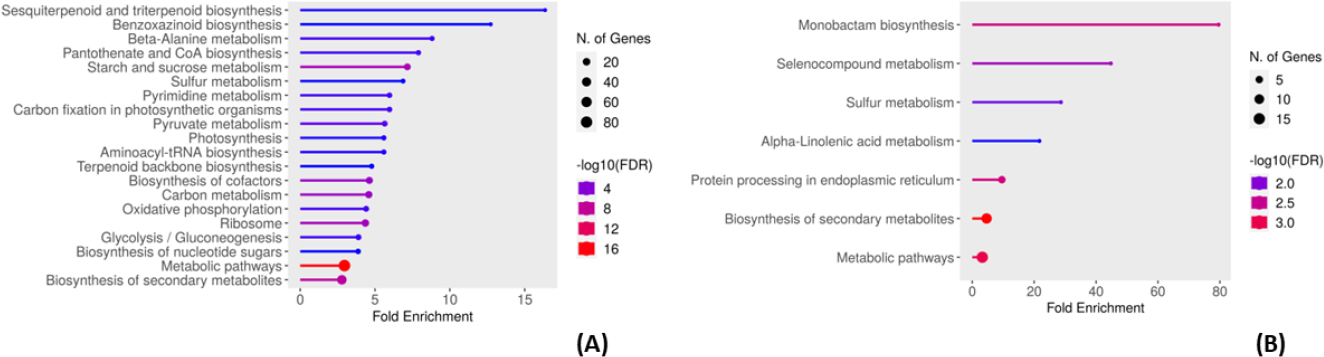
Significantly enriched gene ontology (GO) terms in the **(a)** DM-resistant CML 431 and **(b)** DM-susceptible Pi23

The top GO terms for the DM-resistant CML 431 include ‘Sesquiterpenoid and triterpenoid biosynthesis’, ‘Benzoxazinoid biosynthesis’, etc. On the other hand, only seven (7) GO terms were significantly enriched for the DM-susceptible Pi23. Insights on the genes involved in these biological processes and molecular functions are discussed further.

### Maize pattern recognition receptors (PRR) in response to disease infection

Among the set of DEGs exclusive to the DM-resistant line, we scanned for pattern recognition receptors (PRR) domains which may play roles in response to pathogen infection and recognition. As a result, three (3) leucine-rich repeat (LRR) kinases, one (1) domain of unknown function 26 (DUF26) kinase, one (1) L-lectin kinase, one (1) LysM protein kinase, three (3) receptor-like protein kinases (RLCK), one (1) glycogen synthase kinase (GSK), one (1) casein kinase (CK), two (2) SnRK3 SNF1-related protein kinase, and one (1) calcium-dependent protein kinase (CDPK) were identified. The differential gene expression analysis revealed that three LRRs, with *LRR-VI-2* (zm00001eb295670), *LRR-VII* (zm00001eb177640), and *LRR-XI* (zm00001eb145890), had fold-changes of approximately -2.02-, -1.01-, and -2.3-folds, respectively. Among the RLCK kinases, two were down-regulated (zm00001eb004440 and zm00001eb265280) while one was over-expressed by 3.14 folds (zm00001eb373760; Figure 4). All the DUF26, L-lectin, LysM found were up-regulated while the CK protein kinase (zm00001eb221870) and GSK protein kinase (zm00001eb127490) were found to have downward gene expression levels of ∼-1.8 and ∼-1.4, respectively, upon DM infection in the DM-resistant inbred. Also, the gene expression of CDPK protein kinase (zm00001eb429270) decreased by -2.1-fold. Further, both SnRK3 SNF1-related protein kinases (zm00001eb330600 and zm00001eb312910) were underexpressed by ∼-1.9- and ∼-2.5-folds, respectively. Alternatively, we found two (2) genes that were differentially expressed in the DM-susceptible Pi23 with possible roles in the successful invasion of DM in maize (Figure 5). Specifically, these are acidic chitinase/chitinase16 (zm00001eb301490; down-regulated by ∼-4.05-fold) and 1-phosphatidylinositol-3-phosphate 5-kinase (FAB1; zm00001eb312210); up-regulated by ∼1.56-fold).

**Figure 4.**
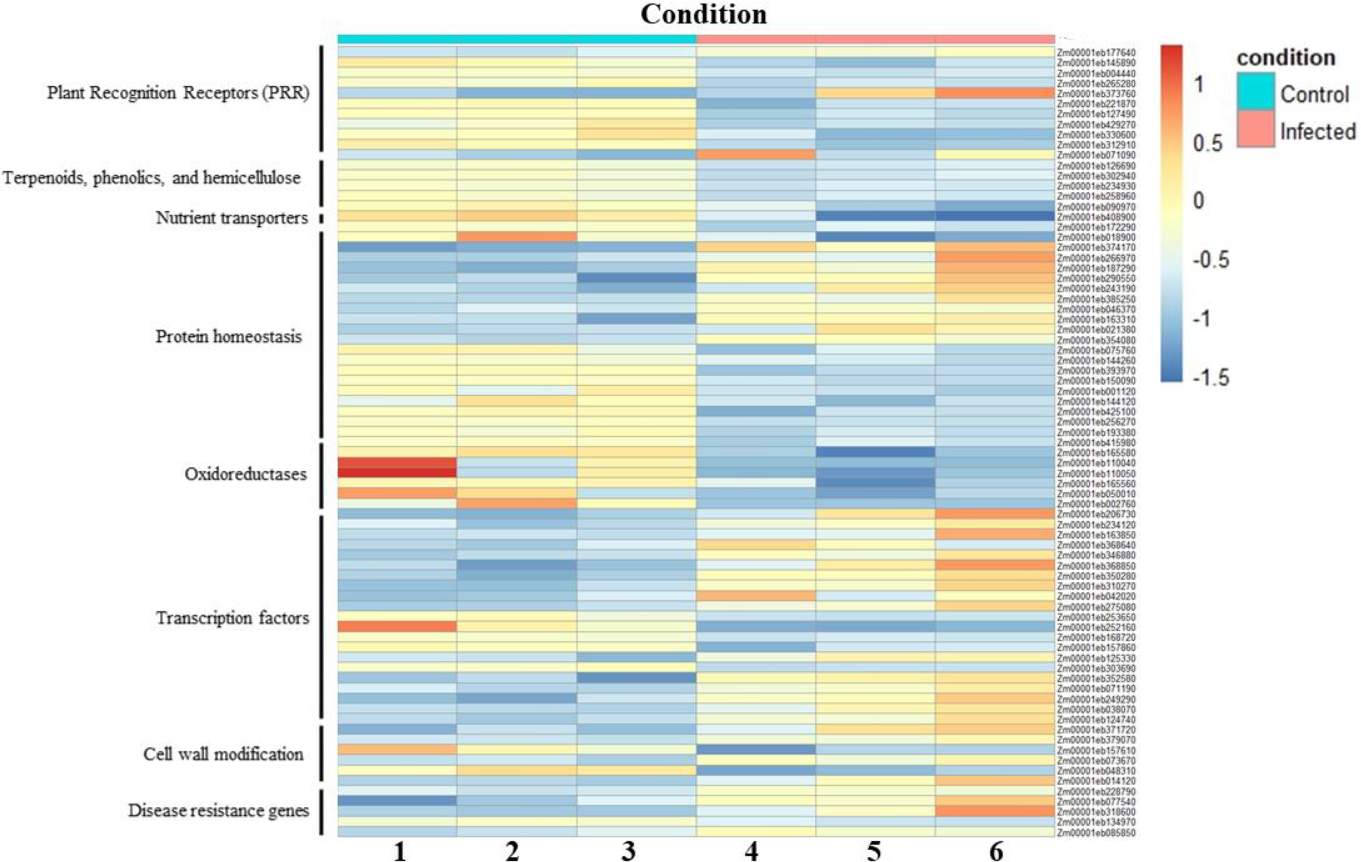
Expression patterns of the DEGs with possible disease-resistance roles found exclusively in the DM-resistant CML 431.

**Figure 5.**
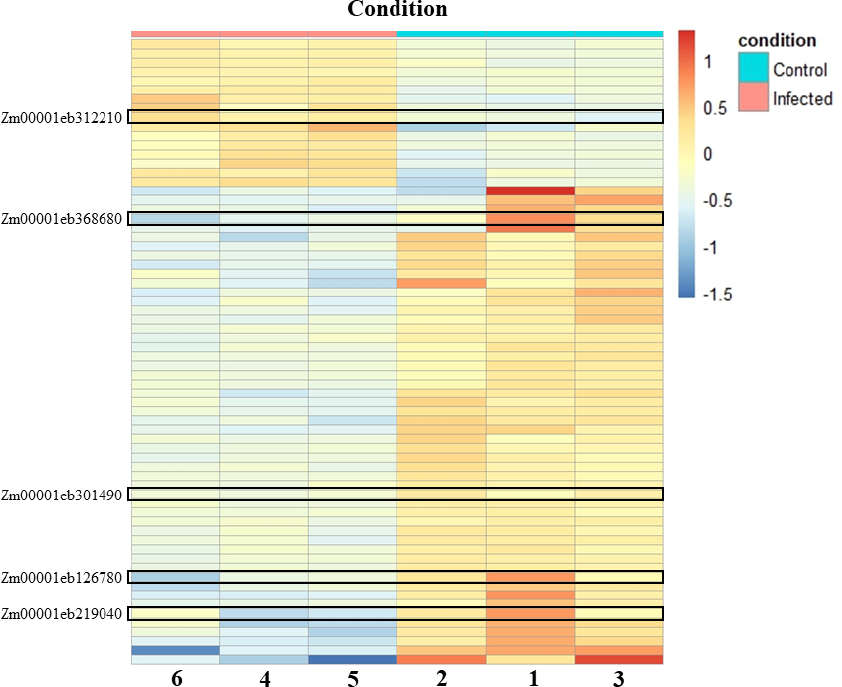
Expression patterns of the 111 DEGs detected in the DM-susceptible Pi23.

### Terpenoid and phenolic biosynthesis genes in response to P. philippinensis invasion

Plants also synthesize secondary and specialized metabolites to defend themselves against pests and diseases. Notably, terpenoids have been identified as key contributors to biotic resistance in maize (Wang et al. 2022; Block et al. 2019). Here, we found five (5) terpenoid biosynthesis genes (Figure 4), all of which were down-regulated except for mono-/sesquiterpene-/diterpene synthase (zm00001eb071090) which was up-regulated by ∼2.36 folds. To list, these include deoxy xylulose reductase 1 (*dxr1*; zm00001eb126690), zeta-carotene desaturase (ZDS; zm00001eb302940), lycopene beta cyclase (LCY-b; zm00001eb234930), and carotenoid beta-ring hydroxylase (LUT5; zm00001eb258960). In a similar case, it was reported that the downregulation of terpene levels, specifically the D-limonene synthase gene, was tightly associated with the increase of innate immune response in oranges against *Penicillium digitatum* infection (Rodrí guez et al. 2014). In contrast, the susceptible maize line, Pi23, has significant down-regulation of genes involved in the biosynthesis of isoflavonoid (*isoflavone reductase-like*; Zm00001eb126780) with a decreased fold-change value of ∼-3.05 (Figure 5).

### Nutrient transporter genes (NTR) in response to pathogen infection in maize

One of the limiting factors during disease infection is the plant’s systemic access to nutrients. Previous studies investigating plant-pathogen interactions have highlighted the significance of nutrient transporter genes (NTRs) in response to plant pathogens (Wang et al. 2022). Specifically, sugar transporter proteins (STPs and SWEETs; Chen et al. 2010) and amino acid transporter proteins (AATs; Sonawala et al. 2018) were reported to be involved in pathogen invasions in crops, including in maize. Here, one AAT and one SWEET were found to be differentially expressed in the DM-resistant line upon inoculation. Amino acid transporter protein zm00001eb090970 was down-regulated by ∼-1.72-fold while *SWEET13B* (zm00001eb408900) was under-expressed by ∼-2.9-fold. Interestingly, an amino acid/auxin permease 57 (*aaap57*) Zm00001eb368680 was exclusively differentially expressed in the susceptible Pi23 genotype, with ∼3.02-fold-change decrease, signifying its possible role in the successful downy mildew progression (Figure 5).

### Homeostasis in resistance response to pathogen invasion

Programmed cell death (PCD) is a mechanism activated by the cell itself to ensure that the correct growth and development occur, while it is also a defense response by plants as a mechanism against environmental stresses, and abiotic, and biotic factors (De Gara and Locato 2018). Two (2) differentially expressed genes involved in PCD were observed in this study. Specifically, genes coding for a protein lysine-specific demethylase (*LSD/LOL*; zm00001eb172290), and a metacaspase-like regulator (*MCP1*; zm00001eb018900) were both down-regulated by ∼-1.18- and ∼-2.78-folds, respectively (Figure 4).

Interestingly, 19 E3 ubiquitin ligase genes involved in protein homeostasis were differentially expressed (Figure 4). Precisely, 10 were up-regulated (zm00001eb374170, zm00001eb266970, zm00001eb187290, zm00001eb290550, zm00001eb243190, zm00001eb385250, zm00001eb046370, zm00001eb163310, zm00001eb021380, zm00001eb354080) while nine (9) were down-regulated (zm00001eb075760, zm00001eb144260, zm00001eb393970, zm00001eb150090, zm00001eb001120, zm00001eb144120, zm00001eb425100, zm00001eb256270, zm00001eb193380) upon DM-infection. Ubiquitination of proteins was previously reported as a key mechanism in plant innate immunity, with E3 ligases as crucial factors in substrate specificity (Marino et al. 2012; Duplan and Rivas 2014).

### Oxidoreductase enzymes in response to pathogen invasion

Cytochrome 450s (CYP) belonging to the oxidoreductase classes of enzymes were also found to be involved in hormone signaling to regulate a plant’s response against external stresses, including disease infections (Li et al. 2010; Pandian et al. 2020). In the DM-resistant yellow maize, we detected seven (7) DEGs encoding for cytochrome 450 enzymes (Figure 4). These include the genes coding for CYP monooxygenase and Trimethyltridecatetraene synthase (zm00001eb415980; down-regulated by ∼-1.7 folds), CYP 71C3 (zm00001eb165580; down-regulated by ∼-2.7 folds), 2 CYP 709B2 (zm00001eb110040 & zm00001eb110050; down-regulated by ∼-8.2 folds and ∼-1.7 folds, respectively), CYP 71a1 & 3-hydroxyindolin-2-one monooxygenase (zm00001eb165560; down-regulated by ∼-1.7 folds), brassinosteroid-deficient dwarf1 (*brd1*; zm00001eb050010; down-regulated by ∼-3.5 folds), and CYP 78A6 (zm00001eb002760; down-regulated by ∼-7.7 folds). Interestingly, the DMR6-LIKE OXYGENASE 2 gene (Zm00001eb219040) was exclusively down-regulated in the DM-susceptible Pi 23 maize inbred line (Figure 5). *DLO1* mutant in Arabidopsis resulted in a degree of resistance to downy mildew disease caused by *Hyaloperonospora arabidopsidis* (Zeilmaker et al. 2015).

### Transcription factors with roles in plant disease resistance and immunity

Ten (10) differentially expressed WRKY transcription factors (TFs) were detected in the DM-resistant maize line (Figure 4). Fascinatingly, all these WRKY TFs were up-regulated for as much as ∼4.7 folds. WRKY factors are involved in the transcriptional reprogramming of the plant’s response toward different invading pathogens (Pandey and Somssich 2009) which can either be positive or negative. Positive influence involves importance in plant defense signaling while negative influence includes their involvement in compromising basal defense and defense gene inactivation. Other DEGs coding for transcription factors with relevant plant defense mechanisms found in the DM-resistant maize in response to DM include 1 B-box (BBX; zm00001eb253650), two (2) MYB-related (zm00001eb252160 & zm00001eb168720), a bZIP TF (zm00001eb157860), seven (7) ethylene response factor (ERF; zm00001eb125330, zm00001eb303690, zm00001eb352580, zm00001eb071190, zm00001eb249290, zm00001eb038070, & zm00001eb124740).

### Cell wall modifications in resistance response to disease pressure

In cereals such as maize, the cell wall is composed of cellulosic microfibrils embedded in a matrix of hemicelluloses and some pectin which have determinative roles in maize pest and disease resistance (Santiago et al. 2013). In our study, one (1) gene involved in hemicellulose biosynthesis was up-regulated while two (2) were down-regulated in the resistant line CML 431 (Figure 4). Specifically, a galacturonosyltransferase gene (zm00001eb371720) was up-regulated by ∼2.7-folds while mannan synthase (CSLD; zm00001eb379070) and endo-beta-1,4-mannanase (zm00001eb157610) were down-regulated. In addition, two (2) pectin biosynthesis, modification, and degradation genes were detected wherein a methyltransferase (QUA; zm00001eb073670) and a pectin methylesterase (zm00001eb048310) were up-regulated by ∼1.7-folds and down-regulated by ∼2.8-folds, respectively. Lastly, a regulatory protein of cellulose-hemicellulose network assembly was found to be down-regulated in the resistant CML 431 (zm00001eb014120).

### Differentially expressed disease-resistance genes in resistant CML 431

Finally, we scanned the Mercator4 results of the exclusive DEGs from the DM-resistant CML 431 inbred line and found five (5) disease-resistance genes or genes coding for disease-resistance proteins (Figure 4). Explicitly, we recognized a pollen-signaling protein1 (*psip1*; zm00001eb228790; up-regulated by ∼0.79-folds) with a disease resistance protein ortholog in Arabidopsis, and two (2) disease resistance proteins namely; putative disease resistance RGA1 (zm00001eb077540; up-regulated by ∼2.09-folds), and disease resistance protein RPM1 (zm00001eb318600; up-regulated by ∼3.36-folds). Furthermore, we also found a NB-ARC domain-containing protein (zm00001eb134970; up-regulated by 0.89-folds) and a RIN4-RPM1 immune signaling regulatory factor (zm00001eb085850; up-regulated by ∼1.48-folds), both involved in the effector-triggered immunity (ETI) machinery defense response in plants.

### Transcriptome-wide variant calling and annotation between DM-infected susceptible and resistant maize

The RNA-seq data generated was also utilized to identify genetic variants that may describe the mechanisms of maize resistance against DM. After consolidation of the three biological replicates per maize line using BCFtools, a total of 30,672 and 29,935 SNPs were mined in the susceptible and resistant maize lines, respectively (Table 1). Conversely, a high amount of indels was found in the resistant line compared to the susceptible. These variants unique between the two would provide insights into the differential response of the susceptible and resistant lines to DM.

**Table 1.** The total number of SNPs and indels detected exclusively between the susceptible and resistant maize.

| Line | SNPs | Indels |
| --- | --- | --- |
| Susceptible (Pi23) | 30,672 | 8,534 |
| Resistant (CML 431) | 29,935 | 134,679 |

**Table 2.** Distribution of the genetic variants and their effects by impact on gene structure.

|  | Pi23 |  | CML 431 |  |
| --- | --- | --- | --- | --- |
| Impact | SNPs | Indels | SNPs | Indels |
| High | 169 | 9,983 | 623 | 23,201 |
| Low | 6,336 | 1,140 | 56,819 | 2,708 |
| Moderate | 4,414 | 481 | 39,001 | 3,324 |
| Modifier | 84,647 | 20,096 | 339,325 | 79,463 |

Variant annotation was performed using SnpEff v.5.1 (Cingolani et al. 2012). The number of effects by impact classified as high, moderate, low, as well as non-coding region/modifier variants was summarized in Table 3. High-impact variants effects are classified as stop gain, stop loss, splice acceptor, and splice donor variants. On the other hand, moderate-impact variants include those that introduce missense mutations while low-impact variants are synonymous (i.e., no change in the amino acid sequence).

**Table 3.** List of resistant-exclusive DEGs with high-impact SNP variants.

| # | Gene ID | Name |
| --- | --- | --- |
| 1 | Zm00001eb064990 | PGR5-like |
| 2 | Zm00001eb003780 | mmp93 |
| 3 | Zm00001eb083840 | php10012 |
| 4 | Zm00001eb325170 | Uncharacterized |
| 5 | Zm00001eb058130 | protein phosphatase homolog121 (prh121) |
| 6 | Zm00001eb193380 | ubiquitin-protein ligase14 (upl14) |
| 7 | Zm00001eb261140 | mate1 – multidrug and toxic compound extrusion1 |
| 8 | Zm00001eb121240 | Uncharacterized |
| 9 | Zm00001eb357290 | Uncharacterized |
| 10 | Zm00001eb423540 | umc2350 |
| 11 | Zm00001eb417320 | TIDP3651 |
| 12 | Zm00001eb265810 | RS2Z35 |
| 13 | Zm00001eb372980 | pco131850 |
| 14 | Zm00001eb299980 | IDP80 |
| 15 | Zm00001eb156600 | pco085330a |

Most of the variants identified were modifiers i.e., intronic variants, UTRs, etc. (25% to 46%, Supplementary Tables S3-S6). Inversely, a high number of high-impact indels were found for both susceptible and resistant maize lines while very few high-impact SNPs were found for both (Table 3). For the resistant maize CML 431, 13.40%, 10.14%, and 8.23% of the indels were splice region variants, splice donor variants, and splice acceptor variants, respectively. Frameshift variants introduced by these indels were at 6.84% while upstream and downstream variants were 6.67% and 11.82%, respectively. The summary of variant effect annotations for the DM-infected susceptible and resistant maize lines was presented in Supplementary Tables S3-S6.

Finally, we scanned for DEGs with high-impact SNPs found in the DM-infected resistant maize line (Figure 12). A total of 15 DEGs were found to contain high-impact SNPs which can potentially be utilized further for DM-resistance breeding in maize (Table 3).

**Figure 12.**
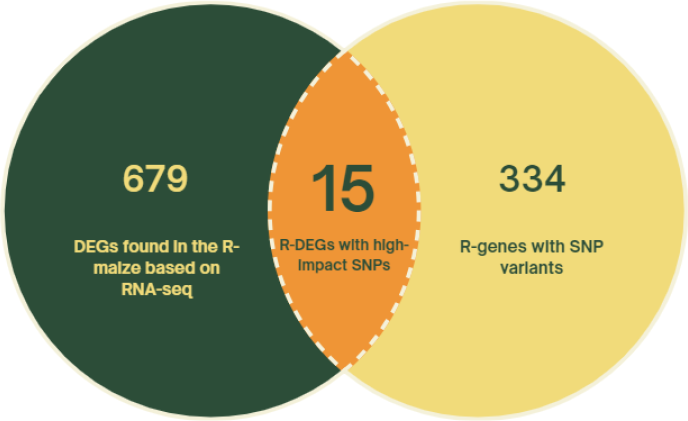
Venn diagram of the resistant-exclusive DEGs and resistant DEGs with SNP variants found in the resistant maize CML 431.

## DISCUSSION

Being one of the most cultivated crops worldwide, maize continues to be plagued by different fungal pathogens such as the southern leaf blight, northern leaf blight, stalk rot, gray leaf spot, and ear rot (Yang et al. 2017). As a consequence, maize developed a complex network of defense systems against pathogen invasion (Wang et al. 2022). Still, only a handful of genes have been studied and validated to confer resistance against disease-causing pathogens in maize. Previously, transcriptomic analyses have been done to elucidate the mechanisms of resistance in maize against different fungal pathogens such as the gray leaf spot (Yu et al. 2018) and ear rot (Yao et al. 2020) but none has been reported yet on *P. philippinensis*, the causal pathogen of the Philippine downy mildew (DM).

In this study, we presented a comparative analysis of maize RNA-seq induced by DM. We identified a total of 111 and 694 DEGs in the DM-susceptible and DM-resistant lines, respectively, of which 43 were shared between the two. PRRs, secondary metabolites, NTRs, transcription factors, protein kinases, etc. were identified to be differentially expressed in the DM-resistant line. This study highlights some of the putative candidate genes as well as gene variants i.e., SNPs and indels, that may be involved or possibly confer maize resistance or susceptibility against DM.

To combat biotic stresses, plants have a two-layered defense system (Jones and Dangl 2006; Zhou and Zhang 2020). In the first layer, pathogen-associated molecular patterns (PAMPs) are recognized by pattern recognition receptors (PRRs) which lead to PAMP-triggered immunity (PTI; Gust et al. 2017). In the second layer, the pathogen effectors are recognized by intracellular nucleotide-binding leucine-rich repeat receptors (NLRs) which activate effector-triggered immunity (ETI; Lolle et al. 2020; Wang et al. 2022). Receptor-like kinases (RLKs) and receptor-like proteins (RLPs) are the major categories of plant PRRs, both of which confer quantitative disease resistance in crops (Kanyuka and Rudd 2019). In this study, we identified 3 LRR kinases,1 L-lectin kinase, 1 LysM protein kinase, 3 RLCKs, 1 GSK, 1 CK, 2 SNF1-related protein kinase, and 1 CDPK that are differentially expressed in the DM-resistant maize. Only a few LRR kinases were reported to be involved in maize disease resistance (Man et al. 2020; Wang et al. 2022), although some of these LRRs in maize have homologs that act as immune receptors found in other crops (Song et al. 2015). For instance, a fungal induced-receptor like protein kinase (FI-RLPK) was found to control the defense response against *Fusarium graminearum* by regulating jasmonic acid and antimicrobial phytoalexin production (Block et al. 2021) while another LRR-RLK was found to confer quantitative susceptibility to maize southern leaf blight (Chen et al. 2023). We also found 1 chitinase (chn16) and 1 phosphatidylinositol 3-phosphate 5-kinase (FAB1) that are differentially expressed in the DM-susceptible Pi23. Chitinase plays an important role in plant resistance against fungal pathogens by degrading chitin, an important component of the cell wall of pathogenic fungi (Xin et al. 2022). It is possible that DM successfully integrated itself in the susceptible line since chn16 was under-expressed and thus was unable to hydrolase DM chitin. On the other hand, host phosphoinositides were found to be recruited by fungal pathogens as susceptibility factors for successful plant invasion. In Arabidopsis for instance, a plasma membrane phosphoinositide PI(4,5)P_2_ was dynamically up-regulated in powdery mildew infection sites while mutants with lower levels of PI(4,5)P_2_ showed inhibited fungal pathogen development, causing resistance (Li et al. 2020). Functional analysis has to be performed on these potential PRRs to investigate their involvement in maize DM resistance and susceptibility Secondary metabolic processes have been well-studied to be elicitors of resistance against an array of biotic factors in several plant species (Block et al. 2019). Here we found several terpenoid biosynthetic pathway genes that were differentially expressed in the resistant maize. In a related study, the resistance to green rot caused by *P. digitatum* was related to the down-regulation of D-limonene in oranges (Rodrí guez et al. 2014). Higher levels of jasmonic and salicylic acids were found upon the down-regulation of this terpenoid gene. This warrants further investigation as to why most of these secondary metabolites were repressed upon DM infection in the resistant line.

Access to nutrients is another limiting factor during pathogen invasion (Wang et al. 2022). Several studies have investigated that nutrient transporter genes are crucial in plant disease resistance (Chen et al. 2010; Moore et al. 2015; Sonawala et al. 2018). Our analysis showed two differentially expressed AAT and SWEET in the DM-resistant line. SWEETs are sugar transporters that are hijacked by pathogens to gain access to sucrose to attain successful invasion (Eom et al. 2019). In this study, SWEET13 was found to be under-expressed by ∼2.9 folds. Mutations in the promoter regions of the SWEET13 ortholog in rice were shown to confer durable resistance against the pathogen *Xanthomonas oryzae* (Zaka et al. 2018). AATs, on the other hand, are membrane-bound transport proteins that facilitate amino acid transfer in plant cells, thereby contributing to plant susceptibility to diseases (Berg et al. 2021; Tu nnermann et al. 2022). Our study showed that amino acid permease (aaap14) was under-expressed in the resistant line. Interestingly, loss-of-function mutations in AAP2 genes in cucumbers were found to contribute robust resistance to *P. cubensis*, the causal pathogen of cucurbit downy mildew (Berg et al. 2021). Conversely, in the DM-susceptible maize line, a significant downregulation of the gene encoding for the amino acid transporter *aaap57* gene was exclusively observed to be under-expressed during *P. philippinensis* attack. Hence, the roles of these genes in DM resistance and/or susceptibility warrant future functional genomics validation (i.e., gene editing for loss-of-function or gene knockout or metabolomics for secondary metabolites).

PCD has a central role in the innate immune response against diseases for both plants and animals (Coll et al. 2011). Two genes involved in PCD namely *LSD/LOL* and *MCP1* were both downregulated in CML 431 upon the onset of DM invasion. A previous study has shown that *LSD1* negatively regulates plant runaway cell death and basal disease resistance (Aviv et al. 2002). Specifically, *LSD1-*mediated cell death and signaling components were related to systemic acquired resistance (SAR), namely salicylic acid (SA)-accumulation and non-expression of pathogenesis-related genes1 (*NIM1/NPR1*). Both SA and *NIM1/NPR1* were known to be associated with many disease-resistance responses and it is possible that *LSD1* antagonizes SA-induced and *NIM/NPR1*-dependent pro-death pathways directly. The expression of various metacaspases in plants has been associated with pathogen invasion (Garcia et al. 2022). For instance, the *LeMCA1* metacaspase, positioned near PCD genes, was found to be upregulated upon the fungal pathogen *Botrytis cinerea* infection in tomato (Hoeberichts et al. 2003). Overexpression of metacaspases, which are possibly related to PCD, was observed in various plant species such as tobacco, pepper, and wheat (Hao et al. 2007; Kim et al. 2013; Wang et al. 2012). Members of *Zea mays* metacaspases (*ZmMCs*) on the other hand, have differential functions in response to different maize diseases (Ma et al. 2021). Thus, further validation of *ZmMCs* functions in maize disease response is necessary in understanding the molecular mechanisms of *MCs* in plant innate immunity. Moreover, 19 DEGs coding for E3 ubiquitin ligases were detected in the resistant maize. Numerous E3 ligase proteins have been identified as plant immunity regulators such as the RING E3 ligase HISTONE MONOUBIQUITINATION1 (*HUB1*) which is involved in resistance against necrotrophic fungal pathogens (Dhawan et al. 2009; Marino et al. 2012). Overexpression of genes coding for E3 ligases was also shown to positively regulate powdery mildew resistance in Chinese wild grapevine, reduce plant susceptibility against *Heterodera schachtii* of Arabidopsis, and enhance tolerance against the fungus *Cladosporium fulvum* in tomato (Yu et al. 2013; Hewezi et al. 2016; Chen et al. 2017). Hence, the differential expression of some of these genes in the DM-resistant maize suggests their potential role in conferring resistance through PCD.

Seven (7) of the 694 DEGs were cytochrome P450 genes, which are associated with plant defense response and many other functions (Liu et al. 2020; Pandian et al. 2020). Recent studies in rice and barley showed that CYP genes were involved in disease resistance suggesting the roles of these 8 DEGs in DM resistance in maize (Ameen et al. 2021; Zhang et al. 2021). In addition, the cytochrome P450 monooxygenase was found to play a role in the partial resistance against *Phytophthora sojae*, the causal pathogen of rot in soybean (Khatri et al. 2022). Their crucial function in the biosynthesis of the benzoxazinoids DIMBOA and DIBOA in grasses has also been established as an induced defense mechanism against invading insect pests and fungal pathogens (Frey et al. 2009). These cytochrome enzymes participate in the catalytic conversion of secondary metabolites such as lignins, terpenoids, phenylpropanoids, alkaloids, and such that are utilized by plants as a response against several biotic and abiotic stresses.

Ten (10) WRKY transcription factors were observed to be overexpressed in DM-resistant maize. Many studies have already shown that WRKYs have pivotal roles in plant defense, while several studies also showed that they can be negative regulators of resistance (Eulgem and Somssich 2007; Pandey and Somssich 2009). For instance, loss-of-function in both *WRKY11* and *WRKY17* increased the basal resistance of *A. thaliana* to *P. syringae* pv tomato (Journot-Catalino et al. 2006) while enhanced resistance was also observed in Arabidopsis in *WRKY7* loss-of-function mutants which were associated with the increased induction of SA-regulated Pathogenesis-related 1 (*PR1*) by the same pathogen (Kim et al. 2006). Transcription factors such as BBX, MYB-related, bZIP, and ERFs, were also found to be differentially expressed in the resistant maize line. The relevance of these transcription factors in plant defense against pathogens has been previously documented in other crops as well (Gangappa and Botto 2014; Sasaki et al. 2015; Froidurea et al. 2010, Wang et al. 2019; Zang et al. 2020; Zhang et al. 2021). Determining the roles and functions of these transcription factors in maize-DM interactions is pivotal in elucidating molecular sources of resistance.

Pathogens encounter the cell wall as the first obstruction in their attempt to invade and colonize the plant (Bacete et al. 2018). Modifications in plant cell walls, either through overexpression or impairment of cell wall-related genes, have profound roles in disease resistance (Bellincampi et al. 2014; Malinovsky et al. 2014). For instance, a reduction in xylan acetylation revealed enhanced resistance in Arabidopsis against the necrotrophic fungus *B. cinerea* and biotroph *H. arabidopsidis* (Manabe et al. 2011; Pawar et al. 2016). In addition, Arabidopsis plants with enhanced levels of wall-bound xylose or with xyloglucan structure modifications have greater resistance to *P. cucumerina* (Delgado-Cerezo et al. 2012). Pectin modification through methylesterification, controlled mainly by pectin methylesterases (PMEs), also has a differential effect on disease response as was found in Arabidopsis and tobacco (Lionetti et al. 2014). Further, these PMEs are post-transcriptionally regulated by endogenous protein inhibitors (PMEIs) which act as mediators of cell wall integrity maintenance relative to plant immunity (Bacete et al. 2018). In our study, several cellulose, hemicellulose, and pectin biosynthesis-associated genes were found to be differentially expressed. Pectin methylesterase26 (*pme26*) was found to be under-expressed by ∼2.8 folds. Analysis of *pme26* activity, pectin methyesterification, as well as the PMEIs modulating this gene, warrants further investigation of DM infection in maize.

Finally, we found a few disease-resistance genes differentially expressed in the DM-resistant CML431 upon DM invasion. Plant NLRs encode immune receptors that interact with pathogen avirulence (*AVR*) gene products, which induce a series of defense responses including hypersensitive response (HR), a type of local PCD (Lai et al. 2020). The *psip1* gene in maize has an ortholog in Arabidopsis (AT4G09435) coding for a disease resistance protein in the NLR protein family. The putative disease resistance protein resistance gene analog1 (RGA1) was also found to be over-expressed in the DM-resistant line by ∼2.09-fold. RGAs including NBS-encoding proteins, RLKs, and RLPs are putative resistance genes with conserved domains and motifs, that are responsible for intracellular signaling and mediation of plant defense genes (Li et al. 2016; Cortaga et al. 2022). Further, two genes coding for disease-resistance proteins At4g33300 and PIK6-NP were also found to be over-expressed, both of which also belong to the LRR protein class. Finally, we found an over-expressed gene coding for RIN4. Previous studies reported the significance of RIN4 in regulating both PAMP- and ETI-triggered immunities in plants (Ray et al. 2019). Hence, these genes may be candidates for downy mildew resistance in maize.

The majority of the genetic variants found for both DM-infected susceptible and resistant maize were modifiers or non-coding variants. Initially, it was thought that intronic variants have little to no effect on gene variants. However, current studies have demonstrated that variations in non-coding regions contribute to different responses in plants. For instance, natural variations in the non-coding region of the NAC-encoding gene *ZmNAC080308* affect the drought tolerance of maize, ultimately improving maize grain yield under drought stress conditions (Wang et al. 2021). Genomic variants are considered high-impact because their presence can have a direct impact on gene functionality (Bohry et al. 2021). High-impact indels typically introduce frameshift mutations while SNPs directly introduce variations within exonic regions. In our study, the majority of the indels found for both susceptible and resistant maize have high-impact effects while most of the SNPs detected have low-to moderate-impact gene effects.

For the eventual marker-assisted breeding of maize with DM resistance, we deliberately shifted our focus on the high-impact SNPs we found in the differentially expressed genes detected in the resistant maize CML 431. A total of 15 DEGs were found to contain potentially gene function-altering SNPs i.e., high-impact SNPs. By unraveling this set of genes and gene variants through transcriptome sequencing-assisted discovery, we can perform targeted genotyping-by-sequencing, single marker association studies, as well as marker-assisted breeding and selection for yellow maize DM resistance.

## CONCLUSION

This study represents the first-ever molecular elucidation of the maize-*P. philippinensis* plant-pathogen interaction, shedding light on the mechanisms underlying successful disease resistance and susceptibility. Through comparative transcriptomics analysis, we have identified putative DM resistance genes in maize, providing an opportunity to design a targeted resistance gene panel for an efficient marker-assisted breeding program focused on improved maize resistance against Philippine downy mildew disease. Additionally, we also identified high-impact gene variants exclusive to both susceptible and resistant maize lines upon DM infection, possibly related to pathogen response and/or resistance.

Our streamlined bioinformatics pipeline revealed a set of important pathogen-response pathways, nutrient transporters, secondary metabolites, regulatory and transcription factors, cell wall modifications, and disease-response genes. Interpretation and further insights into the expression patterns of these genes and proteins would facilitate the mining of putative DM resistance genes that can be seamlessly integrated into breeding programs, thereby assisting in developing DM-resistant maize varieties in the Philippines.

Furthermore, our research unveiled various genes exclusive to the susceptible maize line, suggesting their candidacy as susceptibility factors. These genes can be potential targets for gene editing technologies, such as CRISPR-Cas9, facilitating the development of new lines that could serve as potential parentals for maize hybrid breeding programs. In the long run, this pioneering research lays the essential groundwork for the advancement of a future-proof maize industry, particularly in the face of the widespread prevalence of downy mildew disease, not just in the Philippines, but also on a global scale. By unraveling the genetic factors that influence DM resistance and susceptibility, our findings contribute to the progress of sustainable and resilient maize cultivation practices, ensuring a more secure food supply and strengthening agricultural systems against devastating pathogens.

## ACKNOWLEDGEMENTS

The authors acknowledge the Centro Internacional de Mejoramiento de Maí z y Trigo (CIMMYT), El Batan, Mexico, for the provision of the different CML lines. The authors gratefully acknowledge the Institute of Plant Breeding for the use of its facilities and equipment. We also like to express our gratitude to Ayn Kristina M. Beltran, Rodelio R. Pia, Ranie G. Ramos, Melvin J. Malison, Lorna G. Matanguihan, and Garee R. Hernandez for the technical and administrative assistance. The inputs of Dr. Hayde F. Galvez and Dr. Ma. Anita M. Bautista in initiating the UPLB Basic Research Project are also acknowledged. Our thanks to the Philippine Department of Science and Technology (DOST) ASTI for granting us access to the COARE biocomputing server. We also extend our sincere gratitude for the invaluable assistance and support provided by the Shared Genomics Core Laboratory at the Philippine-California Advanced Research Institute (PCARI), Philippine Genome Center.

## AUTHOR CONTRIBUTIONS

DVL, ANCM, and RRG designed the research. ECJF, EVSA, and JSM performed the experimental setup and preparation of materials. ECJF, ANCM, and DVL conducted the bioinformatics analyses. DVL, ECJF, and ANCM interpreted the results. ECJF and ANCM prepared the graphs. ECJF and DVL wrote the manuscript. DVL secured the project funds. All authors reviewed, revised, and approved the final version of the manuscript.

## DATA AVAILABILITY

The datasets generated during the current study are available from the corresponding author on practical request. Sequence data are available in the NCBI BioProject database (BioProjectID: PRJNA1022320)

## ETHICAL APPROVAL

The authors declare that their work complies with the current laws of the Philippines.

## REFERENCES

Ada EVS, Mendoza JS, Fernandez ECJ, Pia RR, Nun ez JPM, Dela Cueva FM, Maquilan MAD, Manohar ANC, Gardoce RR, Beltran AK, Laude TP, and Lantican DV. 2023. Resistance evaluation of yellow corn (Zea mays L.) inbreds and hybrids against Philippine downy mildew (Peronosclerospora philippinensis Weston Shaw). In: 55th Anniversary and Annual Scientific Conference of the Pest Management Council of the Philippines. July 4-7, 2023.

Aguilar RT, Cortaga CQ, Mendoza JS, Michelmore RM, Martin F and Dela Cueva FM. 2022. Unraveling the genetic structure and diversity of Peronosclerospora philippinensis causing downy mildew disease of sugarcane and corn in the Philippines. Presented during the 2nd IPB-JRO Scientific Conference, University of the Philippines Los Ban os.

Ameen, G. Solanki, S. Sager-Bittara, L. Richards, J. Tamang, P. Friesen, T. L. et al. 2021. Mutations in a barley cytochrome P450 gene enhances pathogen induced programmed cell death and cutin layer instability. PLoS Genet. 17:e1009473. doi: 10.1371/journal.pgen.1009473

Anahosur KH and Patil SH. 1980. Chemical control of sorghum downy mildew in India. Plant Diseases 64: 1004–1006.

Andrews, S. 2010. FastQC: A Quality Control Tool for High Throughput Sequence Data [Online]. Available online at: http://www.bioinformatics.babraham.ac.uk/projects/fastqc/

Aviv DH, Ruste rucci C, Holt BF 3rd, Dietrich RA, Parker JE, Dangl JL. 2002. Runaway cell death, but not basal disease resistance, in lsd1 is SA- and NIM1/NPR1-dependent. Plant J. Feb;29(3):381–91. doi: 10.1046/j.0960-7412.2001.01225.x. PMID: 11844114.

Bacete L, Me lida H, Miedes E, Molina A. 2018. Plant cell wall-mediated immunity: cell wall changes trigger disease resistance responses. Plant J. Feb;93(4):614–636. doi: 10.1111/tpj.13807. Epub 2018 Feb 2. PMID: 29266460.

Bellincampi, D. Cervone, F. and Lionetti, V. 2014. Plant cell wall dynamics and wall-related susceptibility in plant-pathogen interactions. Front. Plant. Sci. 5, 228.

Berg JA, Hermans FWK, Beenders F, Abedinpour H, Vriezen WH, Visser RGF, Bai Y, Schouten HJ. 2021. The amino acid permease (AAP) genes CsAAP2A and SlAAP5A/B are required for oomycete susceptibility in cucumber and tomato. Mol Plant Pathol. Jun;22(6):658–672. doi: 10.1111/mpp.13052. Epub 2021 May 2. PMID: 33934492; PMCID: PMC8126186.

Block, A. K. Vaughan, M. M. Schmelz, E. A. and Christensen, S. A. 2019. Biosynthesis and function of terpenoid defense compounds in maize (Zea mays). Planta 249, 21–30. doi: 10.1007/s00425-018-2999-2

Block, A. K. Tang, H. V. Hopkins, D. Mendoza, J. Solemslie, R. K. du Toit, L. J. et al. 2021. A maize leucine-rich repeat receptor-like protein kinase mediates responses to fungal attack. Planta 254:73. doi: 10.1007/s00425-021-03730-0

Bohry D, Ramos HCC, Dos Santos PHD, Boechat MSB, Are des FAS, Pirovani AAV, Pereira MG. 2021. Discovery of SNPs and InDels in papaya genotypes and its potential for marker assisted selection of fruit quality traits. Sci Rep. 11(1):292. doi: 10.1038/s41598-020-79401-z. PMID: 33431939; PMCID: PMC7801719.

Bolger, A. M. Lohse, M. & Usadel, B. 2014. Trimmomatic: A flexible trimmer for Illumina sequence data. Bioinformatics. doi:10.1093/bioinformatics/btu170.

Bolger, M. Schwacke, R. Usadel, B. 2021. MapMan Visualization of RNA-Seq Data Using Mercator4 Functional Annotations. In: Dobnik, D. Gruden, K. Ramsak, Z. Coll, A. (eds) Solanum tuberosum. Methods in Molecular Biology, vol 2354. Humana, New York, NY. 10.1007/978-1-0716-1609-3_9

Boyd, L. A. Ridout, C. O’Sullivan, D. M. Leach, J. E. and Leung, H. 2013. Plant-pathogen interactions: disease resistance in modern agriculture. Trends Genet. 29, 233–240. doi: 10.1016/j.tig.2012.10.011

Chen, L. Q. Hou, B. H. Lalonde, S. Takanaga, H. Hartung, M. L. Qu, X. Q. et al. 2010. Sugar transporters for intercellular exchange and nutrition of pathogens. Nature 468, 527–532. doi: 10.1038/nature09606

Chen C, Zhao Y, Tabor G, Nian H, Phillips J, Wolters P, Yang Q, Balint-Kurti P. 2023. A leucine-rich repeat receptor kinase gene confers quantitative susceptibility to maize southern leaf blight. New Phytol. May;238(3):1182–1197. doi: 10.1111/nph.18781. Epub 2023 Feb 25. PMID: 36721267.

Chen S, Zhao H, Wang M, Li J, Wang Z, Wang F, Liu A, Ahammed GJ. 2017. Overexpression of E3 Ubiquitin Ligase Gene AdBiL Contributes to Resistance against Chilling Stress and Leaf Mold Disease in Tomato. Front Plant Sci. Jun 30;8:1109. doi: 10.3389/fpls.2017.01109. PMID: 28713400; PMCID: PMC5492635.

CIMMYT Global Maize Program. 2015. “CML1-CML615A_Information.xls”, CIMMYT Maize Lines (CMLs) - Pedigree and characterization data. CIMMYT Research Data & Software Repository Network, V14. Available from https://hdl.handle.net/11529/10246/23.

Cingolani P, Platts A, Wang le L, Coon M, Nguyen T, Wang L, Land SJ, Lu X, Ruden DM. 2012. A program for annotating and predicting the effects of single nucleotide polymorphisms, SnpEff: SNPs in the genome of Drosophila melanogaster strain w1118; iso-2; iso-3. Fly (Austin). 6(2):80–92. doi: 10.4161/fly.19695. PMID: 22728672; PMCID: PMC3679285.

Coll, N. Epple, P. & Dangl, J. 2011. Programmed cell death in the plant immune system. Cell Death Differ 18, 1247–1256. 10.1038/cdd.2011.37

Cortaga, C.Q. Latina, R.A. Habunal, R.R. et al. 2022. Identification and characterization of genome-wide resistance gene analogs (RGAs) of durian (Durio zibethinus L.). J Genet Eng Biotechnol 20, 29. 10.1186/s43141-022-00313-8

Danecek P, Bonfield JK, Liddle J, Marshall J, Ohan V, Pollard MO, Whitwham A, Keane T, McCarthy SA, Davies RM, Li H. 2021. Twelve years of SAMtools and BCFtools. Gigascience. 10(2):giab008. doi: 10.1093/gigascience/giab008. PMID: 33590861; PMCID: PMC7931819.

Dhawan R, Luo H, Foerster AM, Abuqamar S, Du HN, Briggs SD, Mittelsten Scheid O, Mengiste T. 2009. HISTONE MONOUBIQUITINATION1 interacts with a subunit of the mediator complex and regulates defense against necrotrophic fungal pathogens in Arabidopsis. Plant Cell 21: 1000–1019

De Gara L and Locato V. 2018. (eds.) Plant Programmed Cell Death: Methods and Protocols, Methods in Molecular Biology, vol. 1743, 10.1007/978-1-4939-7668-3_1, © Springer Science+Business Media, LLC.

De los Santos WL, Lansigan FP, Hansen J. 2007. Linking Corn Production, Climate Information and Farm-Level Decision-Making: A Case Study in Isabela, Philippines. In: MVK Sivakumar and J Hansen (eds.) Climate Prediction and Agriculture. Springer, Berlin, Heidelberg. 10.1007/978-3-540-44650-7_16.

Dela Cueva, F., Castro AM, and de Torres RL de. 2022. ‘Peronosclerospora philippinensis (Philippine downy mildew of maize)’, CABI Compendium. CABI International. doi: 10.1079/cabicompendium.44646.

Delgado-Cerezo, M. Sanchez-Rodríguez, C. Escudero, V. et al. 2012. Arabidopsis heterotrimeric G-protein regulates cell wall defense and resistance to necrotrophic fungi. Molecular Plant, 5, 98–114.

Dobin, A. et al. 2013. STAR: Ultrafast universal RNA-seq aligner. Bioinformatics 29. 10.1093/bioinformatics/bts635

Duplan and Rivas, 2014. E3 ubiquitin-ligases and their target proteins during the regulation of plant innate immunity. Front. Plant Sci. 5 2014, p. 42

Eom, J. S. Luo, D. Atienza-Grande, G. Yang, J. Ji, C. Thi Luu, V. et al. 2019. Diagnostic kit for rice blight resistance. Nat. Biotechnol. 37, 1372–1379. doi: 10.1038/s41587-019-0268-y

Eulgem T, Somssich IE. Networks of WRKY transcription factors in defense signaling. Curr Opin Plant Biol. 2007 Aug;10(4):366–71. doi: 10.1016/j.pbi.2007.04.020. Epub 2007 Jul 23. PMID: 17644023.

Exconde OR, Raymundo AD, 1974. Yield loss caused by Philippine corn downy mildew. Philippine Agriculturist, 58(3/4):115–120.

Fernandez ECJ, Nun ez JPP, Gardoce RR, Manohar ANC, Bajaro RM, Lantican DV. 2023. Genetic purity and diversity assessment of parental corn inbred lines using SSR markers for Philippine hybrid breeding. SABRAO J. Breed. Genet. 55(3): 598–608. 10.54910/sabrao2023.55.3.1.

Frey M, Schullehner K, Dick R, Fiesselman A, Gierl A. 2009. Benzoxazinoid biosynthesis, a model for evolution of secondary metabolic pathways in plants. Phytochemistry 70(15-16): 1645-1651.

Froidurea, S. et al. 2010. AtsPLA2-α nuclear relocalization by the Arabidopsis transcription factor AtMYB30 leads to repression of the plant defense response. Proc Natl Acad Sci USA 107.

Gangappa, S. N. & Botto, J. F. 2014. The BBX family of plant transcription factors. Trends in Plant Science vol. 19 Preprint at 10.1016/j.tplants.2014.01.010.

Garcia N, Kalicharan RE, Kinch L, Fernandez J. 2023. Regulating Death and Disease: Exploring the Roles of Metacaspases in Plants and Fungi. International Journal of Molecular Sciences. 24(1):312. 10.3390/ijms24010312

Ge SX, Jung D & Yao R. 2020. ShinyGO: a graphical gene-set enrichment tool for animals and plants, Bioinformatics, Volume 36, Issue 8, Pages 2628–2629, 10.1093/bioinformatics/btz931

George, M.L.C. Prasanna, B.M. Rathore, R.S. et al. 2003. Identification of QTLs conferring resistance to downy mildews of maize in Asia. Theor Appl Genet 107, 544–551. 10.1007/s00122-003-1280-6

Ginting C, Prasetyo J, Dirmawati SR, Ivayani, Timotiwu PB, Maryono T, Widyastuti, Chafisa DIR, Asyifa A, Setyowati E, and Pasaribu AHZ. 2020. Identification of Maize Downy Mildew Pathogen in Lampung and the Effects of Varieties and Metalaxyl on Disease Incidence. Annual Research & Review in Biology. 35(7): 23–35.

Gust, A. A. Pruitt, R. and Nu rnberger, T. 2017. Sensing Danger: Key to activating plant immunity. Trends Plant Sci. 22, 779–791. doi: 10.1016/j.tplants.2017.07.005

Hao, L.; Goodwin, P.H.; Hsiang, T. 2007. Expression of a metacaspase gene of Nicotiana benthamiana after inoculation with Colletotrichum destructivum or Pseudomonas syringae pv. tomato, and the effect of silencing the gene on the host response. Plant Cell Rep. 26, 1879–1888.

Hewezi, T. Piya, S. Qi, M. S. Balasubramaniam, M. Rice, J. H. and Baum, T. J. 2016. Arabidopsis miR827 mediates post-transcriptional gene silencing of its ubiquitin E3 ligase target gene in the syncytium of the cyst nematode Heterodera schachtii to enhance susceptibility. Plant J. 88, 179–192. doi: 10.1111/tpj.13238

Hoeberichts, F.A.; ten Have, A.; Woltering, E.J. 2003. A tomato metacaspase gene is upregulated during programmed cell death in Botrytis cinerea-infected leaves. Planta 2003, 217, 517–522.

Jones, J. D. G. and Dangl, J. L. 2006. The plant immune system. Nature 444, 323–329. doi: 10.1038/nature05286

Journot-Catalino N, Somssich IE, Roby D, Kroj T. 2006. The transcription factors WRKY11 and WRKY17 act as negative regulators of basal resistance in Arabidopsis thaliana. Plant Cell. 2006 Nov;18(11):3289–302. doi: 10.1105/tpc.106.044149. Epub. PMID: 17114354; PMCID: PMC1693958.

Kanyuka, K. and Rudd, J. J. 2019. Cell surface immune receptors: The guardians of the plant’s extracellular spaces. Curr. Opin. Plant Biol. 50, 1–8. doi: 10.1016/j.pbi.2019.02.005

Khatri P, Wally O, Rajcan I, Dhaubhadel S. 2022. Comprehensive Analysis of Cytochrome P450 Monooxygenases Reveals Insight Into Their Role in Partial Resistance Against Phytophthora sojae in Soybean. Front Plant Sci. doi: 10.3389/fpls.2022.862314. PMID: 35498648; PMCID: PMC9048032.

Kim, S.-M.; Bae, C.; Oh, S.-K.; Choi, D. 2013. A pepper (Capsicum annuum L.) metacaspase 9 (Camc9) plays a role in pathogen-induced cell death in plants. Mol. Plant Pathol. 14, 557–566.

Kim KC, Fan B, Chen Z. 2006. Pathogen-induced Arabidopsis WRKY7 is a transcriptional repressor and enhances plant susceptibility to Pseudomonas syringae. Plant Physiol. 142(3):1180–92. doi: 10.1104/pp.106.082487. Epub 2006 Sep 8. PMID: 16963526; PMCID: PMC1630724.

Kolde, R. 2012. Pheatmap: pretty heatmaps. R package version, 1(2), 726.

Lai Y, Lu XM, Daron J, Pan S, Wang J, Wang W, Tsuchiya T, Holub E, McDowell JM, Slotkin RK, Le Roch KG, Eulgem T. 2020. The Arabidopsis PHD-finger protein EDM2 has multiple roles in balancing NLR immune receptor gene expression. PLoS Genet. 14;16(9):e1008993. doi: 10.1371/journal.pgen.1008993. PMID: 32925902; PMCID: PMC7529245.

Lantican, D.V., Nocum, J.D.L., Manohar, A.N.C., Mendoza, J.V.S, Gardoce, R.R, Lachica, G.C., Gueco, L.S., Dela Cueva, F.M. 2023. Comparative RNA-seq analysis of resistant and susceptible banana genotypes reveals molecular mechanisms in response to banana bunchy top virus (BBTV) infection. Sci Rep 13, 18719.

Li X. Zhang J.B. Song B. Li H.P. Xu H.Q. Qu B. Dang F.J. Liao Y.C. 2010. Resistance to Fusarium head blight and seedling blight in wheat is associated with activation of a cytochrome P450 gene. Phytopathology. 100:183–191. doi: 10.1094/PHYTO-100-2-0183.

Li, B. & Dewey, C. N. 2011. RSEM: Accurate transcript quantification from RNA-Seq data with or without a reference genome. BMC Bioinformatics. doi:10.1186/1471-2105-12-323.

Li P, Quan X, Jia G et al. 2016. RGAugury: a pipeline for genome-wide prediction of resistance gene analogs (RGAs) in plants. BMC Genomics 17. 10.1186/s12864-016-3197-x

Li Qin, Zhuqing Zhou, Qiang Li, Chun Zhai, Lijiang Liu, Teagen D. Quilichini, Peng Gao, Sharon A. Kessler, Yvon Jaillais, Raju Datla, Gary Peng, Daoquan Xiang, Yangdou Wei. 2020. Specific Recruitment of Phosphoinositide Species to the Plant-Pathogen Interfacial Membrane Underlies Arabidopsis Susceptibility to Fungal Infection, The Plant Cell, Volume 32, Issue 5, Pages 1665–1688, 10.1105/tpc.19.00970

Lionetti, V. Raiola, A. Cervone, F. and Bellincampi, D. 2014. Transgenic expression of pectin methylesterase inhibitors limits tobamovirus spread in tobacco and Arabidopsis. Mol. Plant Pathol. 15, 265–274.

Liu, X. Zhu, X. Wang, H. Liu, T. Cheng, J. and Jiang, H. 2020. Discovery and modification of cytochrome P450 for plant natural products biosynthesis. Synth. Syst. Biotechnol. 5, 187–199. doi: 10.1016/j.synbio.2020.06.008

Lolle, S. Stevens, D. and Coaker, G. 2020. Plant NLR-triggered immunity: From receptor activation to downstream signaling. Curr. Opin. Plant Biol. 62, 99–105. doi: 10.1016/j.coi.2019.12.007

Love, M. I. Huber, W. & Anders, S. 2014. Moderated estimation of fold change and dispersion for RNA-seq data with DESeq2. Genome Biol 15.

Love, M. I. Soneson, C. & Robinson, M. D. 2017. Importing transcript abundance datasets with tximport. dim (txi. inf. rep $ infReps $ sample1) 1.

Lukman R, Afifuddin A, Lubbersted T. 2013. Unraveling the Genetic Diversity of Maize Downy Mildew in Indonesia. J Plant Pathol Microb 4: 162 doi:10.4172/2157-7471.1000162

Ma S, Shi H, Wang GF. 2021. The potential roles of different metacaspases in maize defense response. Plant Signal Behav. 3;16(6):1906574. doi: 10.1080/15592324.2021.1906574. Epub 2021 Apr 12. PMID: 33843433; PMCID: PMC8143262.

Malinovsky, F.G. Fangel, J.U. and Willats, W.G. 2014. The role of the cell wall in plant immunity. Front. Plant. Sci. 5, 178.

Man J, Gallagher JP, Bartlett M. 2020. Structural evolution drives diversification of the large LRR-RLK gene family. New Phytologist 226: 1492–1505.

Manabe, Y. Nafisi, M. Verhertbruggen, Y. et al. 2011. Loss-of-function mutation of REDUCED WALL ACETYLATION2 in Arabidopsis leads to reduced cell wall acetylation and increased resistance to Botrytis cinerea. Plant Physiol. 155, 1068–1078.

Marino D and others 2012. Ubiquitination during Plant Immune Signaling, Plant Physiology, Volume 160, Issue 1, Pages 15–27, 10.1104/pp.112.199281.

Moore, J. W. Herrera-Foessel, S. Lan, C. Schnippenkoetter, W. Ayliffe, M. Huerta-Espino, J. et al. 2015. A recently evolved hexose transporter variant confers resistance to multiple pathogens in wheat. Nat. Genet. 47, 1494–1498. doi: 10.1038/ng.3439

Murray, G. M. 2009. Threat-specific contingency plan: Philippine downy mildew of maize (Peronosclerospora philippinensis) and downy mildew of sorghum (P. sorghi). Australia: Plant Health Australia. https://www.planthealthaustralia.com.au/wp-content/uploads/2013/03/Downy-mildew-of-maize-and-sorghum-CP-2009.pdf

Pandey SP, Somssich IE. 2009. The role of WRKY transcription factors in plant immunity. Plant Physiol. 2009 Aug;150(4):1648–55. doi: 10.1104/pp.109.138990. Epub. PMID: 19420325; PMCID: PMC2719123

Pandian BA, Sathishraj R, Djanaguiraman M, Prasad PVV, Jugulam M. 2020. Role of Cytochrome P450 Enzymes in Plant Stress Response. Antioxidants (Basel). 25;9(5):454. doi: 10.3390/antiox9050454. PMID: 32466087; PMCID: PMC7278705.

Pascual CB, Calilung Jr B, Bituin N, Raymundo AD, Hautea DM, Salazar AM. 2006. Host Resistance and Pathogen Conidial Characteristics Across Locations of Philippine Corn Downy Mildew. The Philippine Agricultural Scientist. 88(4): 489–494.

Pawar, P.M.A. Derba-Maceluch, M. Chong, S.L. et al. 2016. Expression of fungal acetyl xylan esterase in Arabidopsis thaliana improves saccharification of stem lignocellulose. Plant Biotechnol. J. 14, 387–397.

Philippine Statistics Authority (PSA). 2023. 2017-2021 Crops Statistics of the Philippines.

Raudvere, U. et al. 2019. G:Profiler: A web server for functional enrichment analysis and conversions of gene lists (2019 update). Nucleic Acids Res 47.

Ray SK, Macoy DM, Kim WY, Lee SY, Kim MG. Role of RIN4 in Regulating PAMP-Triggered Immunity and Effector-Triggered Immunity: Current Status and Future Perspectives. Mol Cells. 2019 Jul 31;42(7):503–511. doi: 10.14348/molcells.2019.2433. PMID: 31362467; PMCID: PMC6681865.

Raymundo AD, and Exconde OR. 1976. Economic effectiveness of resistant varieties and duter/dithane M-45 foliar spray for the control of Philippine corn downy mildew. Philippine Agriculturist, 60: 52–65.

Rodrí guez A, Shimada T, Cervera M, Alque zar B, Gadea J, Go mez-Cadenas A, De Ollas CJ, Rodrigo MJ, Zacarías L, Pen a L. 2014. Terpene down-regulation triggers defense responses in transgenic orange leading to resistance against fungal pathogens. Plant Physiol. 164(1):321–39. doi: 10.1104/pp.113.224279. Epub 2013 Nov 5. PMID: 24192451; PMCID: PMC3875811.

Ruswandi D, Carpena AL, Lantican RM, Hautea DM, Canama AO, and Raymundo AD. 2014. Genetic Analysis of Components of Resistance and Quantitative Trait Loci Mapping of Philippine Downy Mildew Resistance Gene in Maize (Zea mays L.). Asian Journal of Agricultural Research. 8(3): 136–149.

Salazar AM, Elca CD, Lapin a, and Salazar FJD (Asian Social Project Services Inc.). 2021. Issues Paper on Corn Industry in the Philippines. Philippine Competition Commission (PCC) 01. https://www.phcc.gov.ph/wp-content/uploads/2021/01/PCC-Issues-Paper-2021-01-Issues-Paper-on-Corn-Industry-in-the-Philippines.pdf

Santiago R, Barros-Rios J, Malvar RA. 2013. Impact of cell wall composition on maize resistance to pests and diseases. Int J Mol Sci. 14(4):6960–80. doi: 10.3390/ijms14046960. PMID: 23535334; PMCID: PMC3645672

Sasaki, N. et al. 2015. Transient expression of tobacco BBF1-related Dof proteins, BBF2 and BBF3, upregulates genes involved in virus resistance and pathogen defense. Physiol Mol Plant Pathol 89.

Singh SD, and R. Gopinath. 1985. A seedling inoculation technique for detecting downy mildew resistance in pearl millet. Plant Dis. 69, pp. 582–584

Sonawala, U. Dinkeloo, K. Danna, C. H. McDowell, J. M. and Pilot, G. 2018. Review: Functional linkages between amino acid transporters and plant responses to pathogens. Plant Sci. 277, 79–88. doi: 10.1016/j.plantsci.2018.09.009

Song W, Wang B, Li X, Wei J, Chen L, Zhang D, Zhang W, Li R. 2015. Identification of Immune Related LRR-Containing Genes in Maize (Zea mays L.) by Genome-Wide Sequence Analysis. Int J Genomics. 2015;2015:231358. doi: 10.1155/2015/231358. Epub. PMID: 26609518; PMCID: PMC4645488.

Tunnermann, L. Colou, J. Nasholm, T. and Gratz, R. 2022. To have or not to have: Expression of amino acid transporters during pathogen infection. Plant Mol. Biol. 109, 413–425. doi: 10.1007/s11103-022-01244-1

Van der Auwera GA & O’Connor BD. 2020. Genomics in the Cloud: Using Docker, GATK, and WDL in Terra (1st Edition). O’Reilly Media.

Wang, X.; Wang, X.; Feng, H.; Tang, C.; Bai, P.; Wei, G.; Huang, L.; Kang, Z. 2012. TaMCA4, a novel wheat metacaspase gene functions in programmed cell death induced by the fungal pathogen Puccinia striiformis f. sp. tritici. Mol. Plant Microbe Interact. 25, 755–764.

Wang Y, Li T,Sun Z, Huang X, Yu N, Tai H and Yang Q. 2022. Comparative transcriptome meta-analysis reveals a set of genes involved in the responses to multiple pathogens in maize. Front. Plant Sci. 13:971371. doi: 10.3389/fpls.2022.971371

Wang, L. Xiang, L. Hong, J. Xie, Z. & Li, B. 2019. Genome-wide analysis of bHLH transcription factor family reveals their involvement in biotic and abiotic stress responses in wheat (Triticum aestivum L.). 3 Biotech 9.

Wang, N. Cheng, M. Chen, Y. et al. 2021. Natural variations in the non-coding region of ZmNAC080308 contributes maintaining grain yield under drought stress in maize. BMC Plant Biol 21, 305. 10.1186/s12870-021-03072-9

Woodhouse MR, Cannon EK, Portwood JL, Harper LC, Gardiner JM, Schaeffer ML, Andorf CM. 2021. A pangenomic approach to genome databases using maize as a model system. BMC Plant Biol 21, 385. doi: 10.1186/s12870-021-03173-5.

Xin Y, Wang D, Han S, Li S, Gong N, Fan Y, Ji X. 2021. Characterization of the Chitinase Gene Family in Mulberry (Morus notabilis) and MnChi18 Involved in Resistance to Botrytis cinerea. Genes (Basel). 13(1):98. doi: 10.3390/genes13010098. PMID: 35052438; PMCID: PMC8774697.

Yang, Q. Balint-Kurti, P. and Xu, M. 2017. Quantitative disease resistance: Dissection and adoption in maize. Mol. Plant 10, 402–413. doi: 10.1016/j.molp.2017.02.004

Yao, L. Li, Y. Ma, C. Tong, L. Du, F. and Xu, M. 2020. Combined genome-wide association study and transcriptome analysis reveal candidate genes for resistance to Fusarium ear rot in maize. J. Integr. Plant Biol. 62, 1535–1551. doi: 10.1111/jipb.12911

Yu, Y. Shi, J. Li, X. et al. 2018. Transcriptome analysis reveals the molecular mechanisms of the defense response to gray leaf spot disease in maize. BMC Genomics 19, 742. 10.1186/s12864-018-5072-4

Yu Y, Xu W, Wang J, Wang L, Yao W, Yang Y, Xu Y, Ma F, Du Y, Wang Y. 2013. The Chinese wild grapevine (Vitis pseudoreticulata) E3 ubiquitin ligase Erysiphe necator-induced RING finger protein 1 (EIRP1) activates plant defense responses by inducing proteolysis of the VpWRKY11 transcription factor. New Phytol. 200(3):834–846. doi: 10.1111/nph.12418. Epub 2013 Jul 31. PMID: 23905547.

Zaka A, Grande G, Coronejo T, Quibod IL, Chen CW, Chang SJ, Szurek B, Arif M, Cruz CV, Oliva R. 2018. Natural variations in the promoter of OsSWEET13 and OsSWEET14 expand the range of resistance against Xanthomonas oryzae pv. oryzae. PLoS One. 13(9):e0203711. doi: 10.1371/journal.pone.0203711. PMID: 30212546; PMCID: PMC6136755.

Zang Z, Lv Y, Liu S, Yang W, Ci J, Ren X, Wang Z, Wu H, Ma W, Jiang L and Yang W. 2020. A Novel ERF Transcription Factor, ZmERF105, Positively Regulates Maize Resistance to Exserohilum turcicum. Front. Plant Sci. 11:850. doi: 10.3389/fpls.2020.00850

Zeilmaker, T. Ludwig, N. R. Elberse, J. Seidl, M. F. Berke, L. Van Doorn, A & Van den Ackerveken, G. 2015. DOWNY MILDEW RESISTANT 6 and DMR 6-LIKE OXYGENASE 1 are partially redundant but distinct suppressors of immunity in Arabidopsis. The Plant Journal, 81(2), 210–222.

Zhang M, Liu Y, Li Z, She Z, Chai M, Aslam M, He Q, Huang Y, Chen F, Chen H, Song S, Wang B, Cai H, Qin Y. 2021. The bZIP transcription factor GmbZIP15 facilitates resistance against Sclerotinia sclerotiorum and Phytophthora sojae infection in soybean. iScience. 24(6):102642. doi: 10.1016/j.isci.2021.102642. PMID: 34151234; PMCID: PMC8188564.

Zhang, J. H. Zhao, M. Zhou, Y. J. Xu, Q. F. and Yang, Y. X. 2021. Cytochrome P450 monooxygenases CYP6AY3 and CYP6CW1 regulate rice black-streaked dwarf virus replication in laodelphax striatellus (Falle n). Viruses 13:1576. doi: 10.3390/v13081576

Zhou, J. M. and Zhang, Y. 2020. Plant immunity: Danger perception and signaling. Cell 181, 978–989. doi: 10.1016/j.cell.2020.04.028

